# ProInterVal: Validation of Protein-Protein Interfaces through Learned Interface Representations

**DOI:** 10.1101/2023.12.27.573460

**Authors:** Damla Ovek, Ozlem Keskin, Attila Gursoy

## Abstract

Proteins are vital components of the biological world, serving a multitude of functions. They interact with other molecules through their interfaces and participate in crucial cellular processes. Disruptions to these interactions can have negative effects on the organism, highlighting the importance of studying protein-protein interfaces for developing targeted therapies for diseases. Therefore, the development of a reliable method for investigating protein-protein interactions is of paramount importance. In this research, we present an approach for validating protein-protein interfaces using learned interface representations. The approach involves using a graph-based contrastive autoencoder architecture and a transformer to learn representations of proteinprotein interaction interfaces from unlabeled data, then validating them through learned representations with a graph neural network. Our method achieves an accuracy of 0.91 for the test set, outperforming existing GNN-based methods. We demonstrate the effectiveness of our approach on a benchmark dataset and show that it provides a promising solution for validating protein-protein interfaces.

## Introduction

Proteins, as large and complex molecules, have countless roles in biology. They fulfill essential cell functions by interacting through interfaces, forming complexes that greatly enhance our understanding of biological pathways.^1^ These versatile molecules have diverse functions, including catalyzing chemical reactions, transporting molecules, offering structural support, participating in signaling pathways, and enabling ion transport.^2^ Any abnormal interactions can disrupt their functions, potentially causing detrimental effects on the organism. Thus, the intricate world of proteins and their interactions is vital in understanding and maintaining biological processes.

Investigating protein-protein interactions holds significant importance in disease treatment and pharmaceutical research. Potentially leading to effective treatments, targeted drugs can be developed to disrupt or modulate these interactions by identifying diseasecausing protein-protein interfaces.

Advancements in artificial intelligence (AI) and data accumulation have revolutionized the field of protein-protein interface research. With the rapid growth of genomic and proteomic data, AI algorithms can efficiently analyze large datasets, identify patterns, simulate and predict the three-dimensional structures of protein complexes, and predict protein interfaces. These advancements have accelerated the drug discovery process and enabled researchers to explore a vast landscape of protein-protein interfaces comprehensively.

Proteins are typically represented by their sequence and/or structures. The representation of protein structures may vary as primary (1D), secondary (2D), or tertiary structures (3D) based on the specific context and intended application. Three-dimensional structures capture the arrangement of atoms in a protein and provide valuable information about its folding, functional sites, and interaction surfaces. Deep learning approaches are then employed to analyze and extract meaningful features from these 3D structures. One commonly used approach is convolutional neural networks (CNNs),^3–5^ which can learn hierarchical representations by applying filters to local regions of the protein structure. In CNN-based studies, 3D structures can be converted into 2D contact maps that are translation and rotation invariant. Recurrent neural networks (RNNs)^6^ are also utilized to capture sequential dependencies among residues in the primary sequences of proteins. Additionally, graph neural networks (GNNs) have gained popularity for modeling proteins as graphs, where nodes represent amino acids and edges denote spatial, sequential, and/or functional relationships.^7,8^

Previous studies have predicted protein-protein interfaces using various representations and deep learning approaches.^7–11^ For example, DeepInterface^10^ is a CNN-based method that takes two proteins as input and predicts whether they form a biologically valid complex interface. It utilizes voxelized representations of proteins. DOVE^9^ is another CNN model which examines the protein-protein interfaces using 3D voxels, taking into account atomic interaction types and their energetic contributions as input characteristics applied to the neural network. However, CNNs are not rotation invariant, requiring data augmentation to mitigate orientation sensitivity, which affects computational performance during training.

Alternatively, representing protein-protein interfaces as rotation-invariant graphs enables the use of GNNs to overcome these limitations. GNNs handle rotation-invariant graph representations without extensive data augmentation. GNN-DOVE^8^ employs a graph neural network model, representing atoms’ chemical properties as node features and inter-atom distances as edge features. Similarly, DeepRank-GNN^7^ is a GNN-based approach that scores docking models by converting interface residues into two graphs: one representing internal edges between residues from the same chain and another for external edges between residues from different chains. These graphs are then sequentially processed by a dedicated graph interaction neural network.

With the advancements in graph-based deep learning methods, the use of graph-based protein representation learning has gained considerable attention due to its ability to capture the complex interactions and structural dependencies in protein-protein interfaces. However, there is still a need for improvement in the accuracy of the existing methods. One problem to overcome is that handcrafted features limit the expressivity of protein-protein interface representations.

Notably, a recent study^12^ explores the power of learning protein representations based on their 3D structures. Leveraging multiview contrastive learning and self-prediction tasks, essential geometric features of proteins are effectively captured.

The power of contrastive learning is also showed by another study, called ConPLex.^13^ It is a deep learning model that leverages pretrained protein language models (PLex) and employs a protein-anchored contrastive co-embedding (Con) to predict drug-target interactions. After that, a different study has integrated pretrained protein language models into geometric deep learning networks to enhance protein representation learning.^14^

Furthermore, the potential of the use of transformers to investigate protein-protein interfaces has been demonstrated by a recent approach, called PIsToN.^15^ This approach differentiates native-like protein complexes from incorrect conformations by transforming protein interfaces into two-dimensional interface maps. It combines Vision Transformer^16^ architecture with hybrid and multi-attention networks.

This study presents a graph-based approach, called ProInterVal, for validating the biological relevance of protein-protein interfaces using learned interface representations. To validate biological relevance, our model distinguishes between positive and negative interfaces. In the context of our work, a positive interface refers to the interface found in a protein-protein complex that exists in Protein Data Bank (PDB)—meaning it is biologically relevant. Conversely, a negative interface corresponds to the interface observed in a docked protein-protein complex with an unacceptable Critical Assessment of Predicted Interactions (CAPRI) score—meaning it has a fraction of native contacts (Fnat) value lower than 0.1, ligand root mean square deviation (LRMSD) greater than 10Å, and interface root mean square deviation (iRMSD) greater than 4Å. Our approach involves using a novel graphbased contrastive autoencoder architecture and a transformer to learn representations of protein-protein interaction interfaces from unlabeled data, then validating them through learned representations with a graph neural network. We use a few handcrafted features for the generation of the input graph to the representation learning part. On the other hand, the input to the protein-protein interface validation part is the learned representation of the input graph. Therefore, no handcrafted features are used for validating interfaces.

The generation of adequate representations for protein-protein interfaces is accomplished through the utilization of a graph autoencoder. To enhance the representation quality, a transformer component is incorporated to predict edge features. Furthermore, as in a previous study,^12^ our model incorporates a contrastive layer to capture the similarity between correlated substructures within proteins and an edge message passing layer to capture the interdependence among different interactions involving a residue and its neighboring residues. However, our method has the following differences: they apply contrastive autoencoder to the graph of the whole proteins, but we extract and encode only the graph of the interface regions of the protein complexes; the task is different such that they used learned representations to identify protein function and fold classification, whereas we used them to validate proteinprotein interfaces in the docking complexes. Moreover, our model involves a transformer component they do not have and a GCL-based autoencoder different from their autoencoder architecture.

Notably, ProInterVal distinguishes itself from other compared models by capitalizing on learned representations of protein-protein interfaces. Our method exhibits a test set accuracy of 0.91, surpassing the performance of current GNN-based approaches. These results highlight the power of learned representations in accurately assessing protein-protein interfaces. Moreover, the versatility of the learned representations extends to various downstream tasks. For example, they can be utilized for predicting Gene Ontology (GO) terms, providing valuable insights into the functional annotations of proteins. Additionally, these representations enable the classification of protein complexes as either crystal or biological.

## Data & Methods

In this section, we delve into the architecture of our model, which combines a graphbased contrastive autoencoder with a transformer for learning an adequate representation of protein-protein interfaces. The input to our model is a protein complex comprising two chains. Initially, we extract the interface and convert the interface structure into a graph representation. We then process this graph using a graph-based contrastive autoencoder combined with a transformer architecture to acquire a latent representation of the graph. Finally, based on the learned representation, we utilize a graph neural network (GNN) model to generate an interaction score for the two chains. Through this multi-step process, our model effectively validates protein-protein interactions.

### Dataset for Interface Representation Learning

We employed multiple data sources for training and testing our model. We utilized a large set of unlabeled protein-protein complexes for training and testing our graph-based contrastive autoencoder combined with a transformer for generating protein representations. These complexes were obtained from the Protein Data Bank (PDB).^17^ Sequential and structural comparison techniques are employed to create a unique set.^18^ MM-align^19^ is used to compare the complete interface structures (interface structure with both chains) and TM-align^20^ is used to compare one side (single chain) of the interfaces. The dataset is comprised of a total of 534,203 samples.

By leveraging this extensive dataset, we aimed to capture a diverse range of proteinprotein interactions and their corresponding interfaces. The availability of a sizable unlabeled dataset allowed our model to learn robust representations without being limited by labeled data. In the subsequent sections, we describe how this data was processed and used in training our graph-based framework for investigating protein-protein interfaces.

### Dataset for Protein-Protein Interface Validation

#### DeepInterface Dataset

The DeepInterface dataset^10^ served as a crucial component for the validation of proteinprotein interactions. This dataset was obtained from four sources: DOCKGROUND,^21^ PPI4DOCK,^22^ PIFACE,^23^ and PDB.^17^

Positive interfaces were primarily obtained from PIFACE, which contains interface structures extracted from PDB files published until 2012. Additional positive interfaces were extracted from PDB files deposited after 2012 to avoid redundancy. Negative interfaces were derived from DOCKGROUND and PPI4DOCK decoy sets, using CAPRI’s classification.^24^ Incorrect complexes have Fnat (fraction of native contacts) value lower than 0.1, LRMSD (ligand root mean square deviation) greater than 10 Å, and iRMSD (interface root mean square deviation) greater than 4 Å.

Table 1 showcases the quantitative distribution of samples from each source, emphasizing their contribution to the dataset and the partitioning scheme into distinct subsets for training, validation, and test purposes.

**Table 1:**
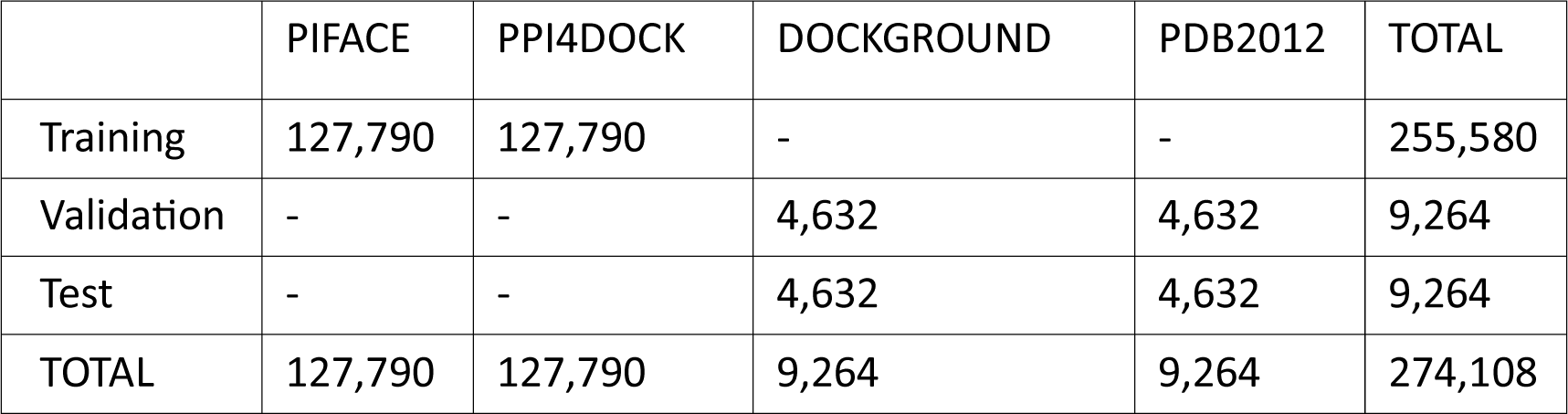
Distribution of positive and negative protein interface data from diverse sources within the DeepInterface dataset, illustrating the partitioning scheme into distinct subsets for training, validation, and test purposes.

|  | PIFACE | PPI4DOCK | DOCKGROUND | PDB2012 | TOTAL |
| --- | --- | --- | --- | --- | --- |
| Training | 127,790 | 127,790 | - | - | 255,580 |
| Validation | - | - | 4,632 | 4,632 | 9,264 |
| Test | - | - | 4,632 | 4,632 | 9,264 |
| TOTAL | 127,790 | 127,790 | 9,264 | 9,264 | 274,108 |

To ensure a fair comparison, we evaluated our model alongside two graph-based docking model evaluation models, namely GNN-DOVE and DeepRank-GNN. The evaluation was performed on three datasets: DeepInterface dataset, GNN-DOVE dataset, and DeepRankGNN dataset.

#### GNN-DOVE Dataset

GNN-DOVE utilizes a combination of the Dockground dataset^21^ and ZDOCK,^25^ as well as a CAPRI scoring dataset.^26^ To eliminate redundancy, the complexes were grouped based on sequence alignment and TM-align.^27^ Specifically, two complexes were assigned to the same group if any pair of proteins from the two complexes exhibited a TM-score greater than 0.5 and a sequence identity of 30% or higher. Dockground and ZRANK are used for training and validation, while the CAPRI scoring dataset is designated for testing purposes.

#### DeepRank-GNN Dataset

DeepRank-GNN dataset is composed of the Docking Benchmark version 5 (BM5).^28^ The BM5 comprises a non-redundant set of 142 complexes (dimers), excluding antibody-antigen complexes and complexes with more than two chains. A total of 25,300 models per complex were generated using the integrative modeling software HADDOCK.^29^ The test set consisted of all docking models generated for 15 randomly selected complexes (379,500 models, 10% of the dataset), while the remaining 127 complexes were split into a training set (80%, 102 complexes) and a validation set (20%, 25 complexes).

### Protein-Protein Interface Representation Learning

#### Input Preparation & Graph Construction

In our study, the initial step involves the extraction of the binding region within a protein complexes. To achieve this, we employ a procedure to identify the residues that interact with each other. Precisely, we assess the proximity of residues from different chains by comparing their distances concerning the sum of their van der Waals radii and an additional threshold of 0.5 Å. If the distance between two residues satisfies this criterion, we consider them contacting residues. Once all the contacting residues are identified, we define neighboring residues. Neighboring residues are determined as those within a distance of 6 Å from any contacting residue within the same chain. Consequently, the union of the contacting and neighboring residues defines the interface region of the protein complex.

Subsequently, we represent the interface region as a graph, wherein residues are represented as nodes, and their interactions are captured through edges. Each node is encoded as a vector of length 30, encompassing the features: residue type, polarity, residue charge, relative accessible surface area (relASA) in the monomer and in the complex form, knowledge-based pair potential (PP), and backbone dihedral angles (*ϕ* and *ψ*). Naccess is used to calculate relASA,^30^ and knowledge-based PP are obtained from Keskin *et. al.*^31^

Naccess has various hyperparameters such as z-slices, probe size, Van der Waals values file, and parameters, whether to ignore HETATM records, hydrogens, and waters.^30^ In this study, we have used default parameter values while running Naccess.

As in previous research of Zang *et. al.*,^12^ we use three distinct edge types in our graph representation:

- Sequential edges: We establish an edge between the *i^th^* and *j^th^* residues if their sequential distance, denoted as |*j* − *i*|, falls below a predefined threshold. Specifically, an edge is created when |*j* − *i*| *<* 3, signifying proximity within the sequence.
- Radius edges: Besides sequential edges, we introduce edges between two nodes, i and j, if the Euclidean distance between them is less than a specified threshold of 10 Å. This criterion ensures that residues in close physical proximity are interconnected.
- K-nearest neighbor edges: Recognizing the potential variations in spatial scales across different proteins, we incorporate edges by connecting each node to its k-nearest neighbors based on the Euclidean distance. We set *k* = 10. This strategy ensures a comparable density of spatial edges across diverse protein graphs, facilitating meaningful graph representations.

The size of the graph and, hence, the adjacency matrix vary according to the size of the input protein-protein interface. However, each node and each edge are represented with fixed-size vectors of length 30 and 3, respectively. The embedding dimension is, on the other hand, fixed to 128×512.

By employing these strategies, our graph-based approach enables comprehensive and informative encoding of the interface region, capturing sequential and spatial relationships between residues within the protein complex. Figure 1 shows the 3D structure and the sequence of a protein-protein complex, interface residues are highlighted, and its graph representation. The nodes are represented as tensors, encompassing residue-specific features, while the edges are encoded as vectors of 1s and 0s, indicating the presence or absence of specific edge types.

**Figure 1:**
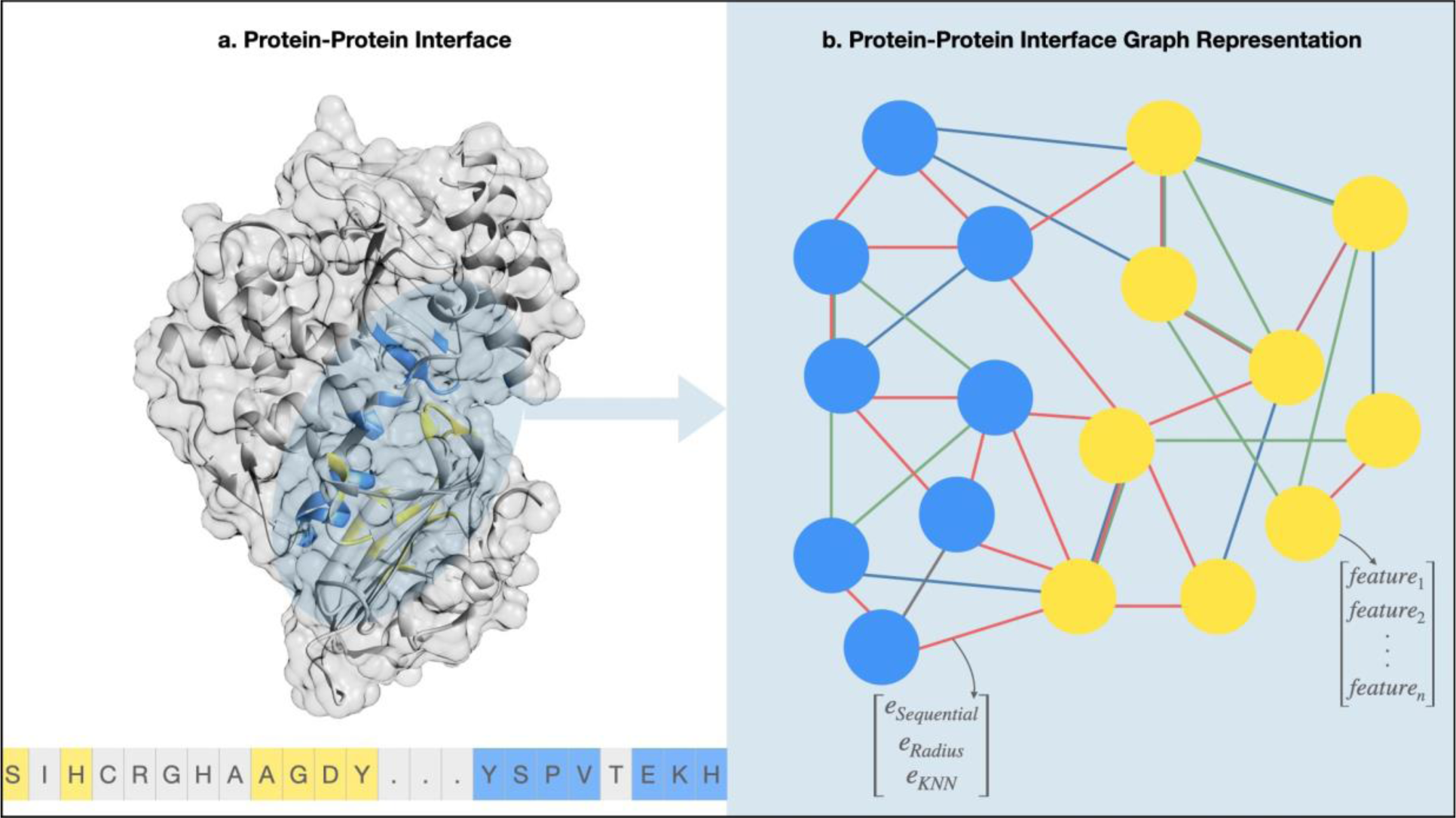
Protein-protein interface structure, sequence, and graph representation. (a) Visualization of a protein-protein complex (PDB ID: 1JTG_A_B) in three-dimensional (3D) structure, depicted as a surface and ribbon model. The sequence of the complex is displayed at the bottom, with interface residues highlighted in color. Residues from chain A are shown in blue, while residues from chain B are depicted in yellow. (b) The corresponding graph representation of the complex is presented. The graph consists of nodes, each representing a residue and its associated features, and edges, representing different types of interactions. Red: Radius Edges, Green: Sequence Edges, Blue: KNN Edges.

#### Graph-based Contrastive Autoencoder

After the construction of the graph, it is passed through a novel graph autoencoder architecture combined with a transformer module to obtain a latent representation. Our proposed model is visually depicted in Figure 2.a. The encoder component inputs the node feature matrix *X* and the adjacency feature tensor *A*, effectively encoding them into a continuous latent representation denoted as *z*. Conversely, given a point in the latent space, the decoder generates an output node feature matrix *X*^′^. Additionally, utilizing a disentangled latent space, the model predicts graph-level properties, yielding the graph property prediction denoted as *y*^′^.

**Figure 2:**
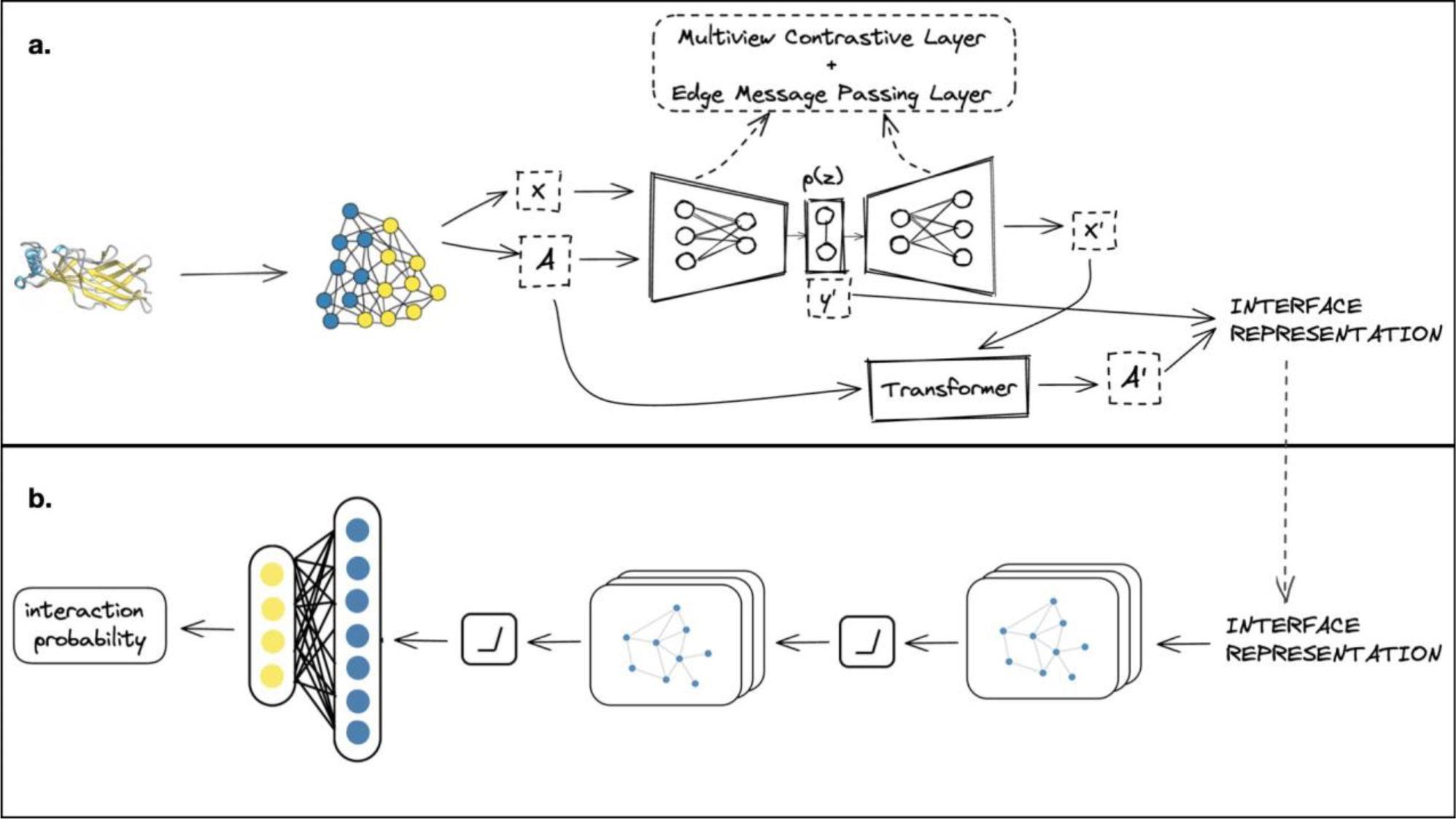
Overall framework. (a) The protein-protein interface structure is represented as a residue-level graph. The graph is inputted into a graph-based autoencoder combined with a transformer to learn the interface representation. (b) The learned representation is then passed through a GNN model to obtain the interaction probability.

#### Multiview Contrastive Layer

A multiview contrastive learning layer was added to the autoencoder to enhance the model’s performance. We employed the same approach taken by Zhang *et. al.*^12^ The multiview contrastive learning involved the application of two cropping functions, namely subsequence, and subspace.

1. A consecutive protein segment was randomly sampled in the subsequence cropping, and the corresponding subgraph was extracted.
2. In the subspace cropping, a center residue was first sampled, and then all residues within a specified distance threshold were selected to form the subgraph.

Furthermore, a random transformation function was applied to the output subgraph, with options including identity (no transformation) and random edge masking (random removal of a fixed ratio of edges from the graph).

#### Edge Message Passing Layer

We added a message-passing layer to enhance the model performance further, as in a prior study.^12^ In message passing, a relational graph, commonly called a line graph, establishes connections among edges. Each node in the line graph represents an edge from the original graph. Specifically, an edge (*i,j,r*_1_) in the original graph is linked to an edge (*w,k,r*_2_) in the line graph if and only if *j* = *w* and *i*/= *k*. Subsequently, a relational graph convolutional network is applied to the line graph, enabling the derivation of the message function for each edge. The message function of an edge is updated by aggregating features from its adjacent edges.

#### Transformer

The proposed model incorporates a transformer component for edge prediction in graphbased architectures inspired by the study of J. Mitton *et al*.^32^ The transformer takes the adjacency feature tensor and the predicted node feature matrix as input and generates a predicted edge feature tensor. The utilized transformer architecture follows a vanilla implementation.^33^

The vanilla transformer architecture is a neural network framework initially designed for natural language processing tasks. At its core, it relies on a multi-head self-attention mechanism, which allows the model to dynamically weigh the importance of different input elements. It consists of an encoder-decoder structure, with both components comprising identical layers. Each layer contains two main sub-layers: a multi-head self-attention mechanism and a position-wise feedforward neural network that processes the attended representations, thus enabling the transformer to capture complex relationships and patterns in data.

During the generation phase, the transformer sequentially adds edges to the graph. At each step, the input to the transformer consists of the node encoding of adjacent nodes to the predicted edge and the previously predicted edges in the graph. This iterative process allows the model to progressively construct the graph by making informed edge predictions based on the available information.

The model effectively captures dependencies and interactions between nodes in the graph by employing a transformer-based approach, enabling accurate edge predictions. The iterative generation of edges ensures that the model incorporates the evolving graph structure while maintaining consistency with the existing edges in the graph.

The objective function of the transformer model can be expressed as:

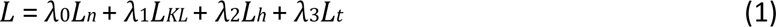

where *λ_i_* represents scaling constants with the values *λ*_0_ = 0.2, *λ*_1_ = 0.3, *λ*_2_ = 0.3, and *λ*_3_ = 0.2; *L_n_* denotes the cross entropy loss for reconstructing node features, *L_KL_* corresponds to the KL divergence loss to calculate the statistical distance between the input graph and the latent space representation, *L_h_* represents the mean squared error loss for predicting graph properties from the latent space, and *L_t_* represents the cross entropy loss for reconstructing edge features.

To optimize the model, an Adam optimizer is employed with an initial learning rate of 5_e−4_.

### Protein Function Analysis of Learned Representations

t-SNE is a technique used for reducing the dimensionality of data in order to visualize the similarities among high-dimensional data points. To analyze the learned protein interface representations t-SNE was employed to project them onto a two-dimensional (2D) space. A random sample set of 10,000 samples was selected, representing five function classes: antibodies, enzymes, receptors, hormones, and transporters. Within this sample set, there were 1,879 antibodies, 849 enzymes, 1,572 receptors, 2,012 hormones, and 3,688 transporters.

A t-SNE plot was generated to visualize the clustering patterns of these classes.

### Protein-Protein Interface Validation

Once the learned representation of two given proteins is obtained, a graph neural network model is employed to predict the interaction score between the given proteins. The GNN model consists of graph convolution layers (GCL), non-linear activation functions (in this case, Rectified Linear Units or ReLU), and pooling layers. The last layer of the model is fully-connected. This architectural design enables the model to capture and process the structural information encoded in the graph representation. The output of this GNN model is a probability indicating the likelihood of the given protein complex being a biologically valid interaction.

The cross-entropy loss was used as the loss function to train the model, and the Adam optimizer was used for parameter optimization. The model was trained for 20 epochs with a batch size of 32.

## Results and Discussion

### Ablation Study

To assess the individual contributions of specific components in our model, we conducted ablation experiments on the PPI validation task. We systematically remove key elements, namely the multiview contrastive layer and the edge message passing layer. Then, we trained each model with the DeepInterface training set and compare prediction accuracy on the test set. This rigorous analysis allowed us to evaluate the impact of each component on the overall performance of the model and gain insights into their respective roles in the learning process. As shown in Table 2, the results can be significantly improved by using multiview contrastive learning and the prediction performance further increases after performing edge message passing.

**Table 2:** Ablation experiment results of graph autoencoder, multiview contrastive layer, and edge message passing layer on the PPI validation task.

| Model | Accuracy |
| --- | --- |
| Graph autoencoder | 0.764 |
| Multiview contrastive graph autoencoder | 0.888 |
| Graph autoencoder with edge message passing | 0.879 |
| ProInterVal | 0.923 |

Multiview contrastive layer is designed to capture the similarity between correlated protein substructures by constructing informative views of the substructures. The layer’s objective is to preserve the similarity between these substructures both before and after mapping them to a latent space. By creating multiple views that reflect different aspects of the protein substructures, the multiview contrastive layer enables the model to capture and preserve the intricate relationships and similarities among the substructures. This contributes to the overall effectiveness of the model in capturing the complex nature of protein interactions and representations.

The edge message passing layer can be regarded as a modified version of the pair representation update specifically tailored for graph neural networks. Drawing inspiration from AlphaFold2,^34^ which utilizes the triangle attention mechanism in transformers to capture pair representations, our edge message passing layer aims to capture the interdependence among various interactions involving a residue and its sequentially or spatially adjacent residues. Unlike the triangle attention in AlphaFold2, our approach incorporates angular information to effectively model diverse types of edge interactions, enabling more efficient sparse edge message passing operations.

### PPI Validation Performance Evaluation

We conducted a comprehensive performance comparison of our ProInterVal, with the two state-of-the-art methods, GNN-DOVE, and DeepRank-GNN, for validating protein-protein interactions. To ensure a fair evaluation, all models were trained from scratch on three distinct training sets and evaluated on their respective test sets. The details of these sets are explained in the dataset section. Notably, the analysis revealed that ProInterVal consistently outperformed the other models across all three datasets. Figure 3 visually depicts the performance of ProInterVal compared to GNN-DOVE and DeepRank-GNN on these datasets. These findings highlight the robustness and efficacy of the ProInterVal model in accurately validating PPIs, underscoring its potential as a powerful tool in computational protein studies.Recently a study based on 2D images of proteins has been published.^15^ Different from our method, it is not a graph-based method, and has ROC-AUC score of 0.81 on the CAPRI score dataset, while our method’s ROC-AUC score on the DeepInterface dataset, constructed based on CAPRI classification, and the GNN-DOVE dataset, sourced from the CAPRI score dataset is 0.88.

**Figure 3:**
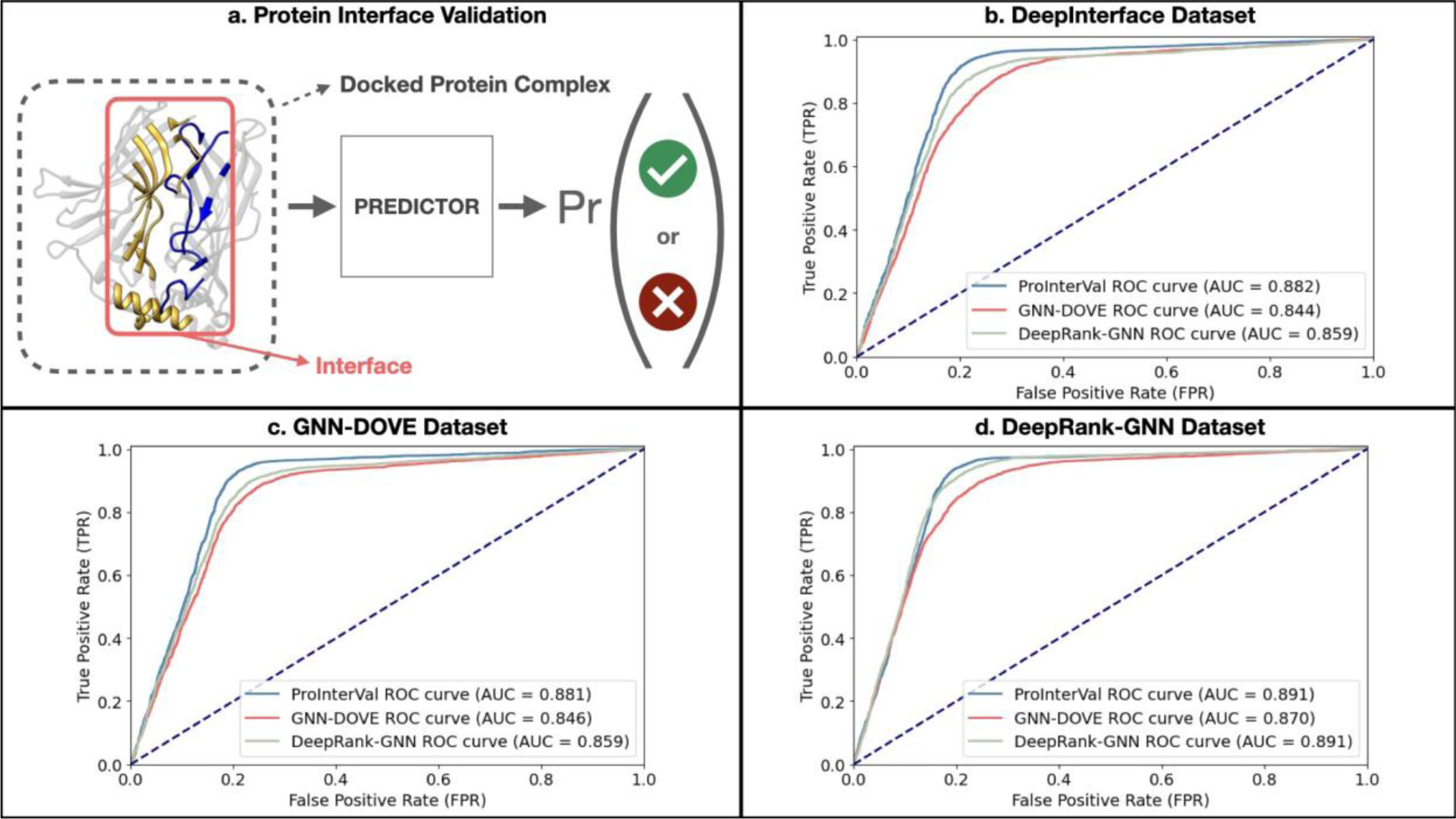
Protein-protein interface validation and performance comparison of ProInterVal, GNN-DOVE, and DeepRank-GNN on multiple datasets. (a) Illustration depicting the concept of protein-protein interface validation, where the predictor generates a probability for the docked complex interface to be a biological complex. (b) Performance evaluation results of ProInterVal, GNN-DOVE, and DeepRank-GNN on the DeepInterface test set, after retraining the models from scratch on the training set of DeepInterface dataset. (c) Performance evaluation results of ProInterVal, GNN-DOVE, and DeepRank-GNN on the GNNDOVE test set, after retraining the models from scratch on the training set of GNN-DOVE dataset. (d) Performance evaluation results of ProInterVal, GNN-DOVE, and DeepRankGNN on the DeepRank-GNN test set, after retraining the models from scratch on the training set of DeepRank-GNN dataset.

ProInterVal distinguishes itself from the compared models by leveraging learned representations of protein-protein interfaces, whereas the other models rely on handcrafted features to represent these interfaces. Protein function is governed by a wide range of structural motifs, characterized by intricate spatio-chemical arrangements of atoms and amino acids. Handcrafted features struggle to comprehensively capture these patterns. Comparative modeling faces challenges in detecting these motifs, as the set of sequence/conformational perturbations that preserve function remains unknown. Therefore, this study introduces an approach that utilizes learned interface representations for validating protein-protein interfaces. Furthermore, as outlined in the methodology section, our model is trained on a dataset of over 500,000 interfaces, enabling it to capture diverse information and exhibit generalizability. t-SNE Analysis

Figure 4 showcases the t-SNE visualizations of protein function clusters and positive/negative interfaces, offering valuable insights into the arrangement of protein functions and the differentiation between positive and negative interfaces. These visualizations demonstrate the potential of our approach in comprehending protein interactions and verifying their biological significance. As depicted in Figure 4.a, the proteins exhibit clustering based on their functional classes in the embedding space, indicating that our model has captured functional fingerprints of the proteins to some extent.

**Figure 4:**
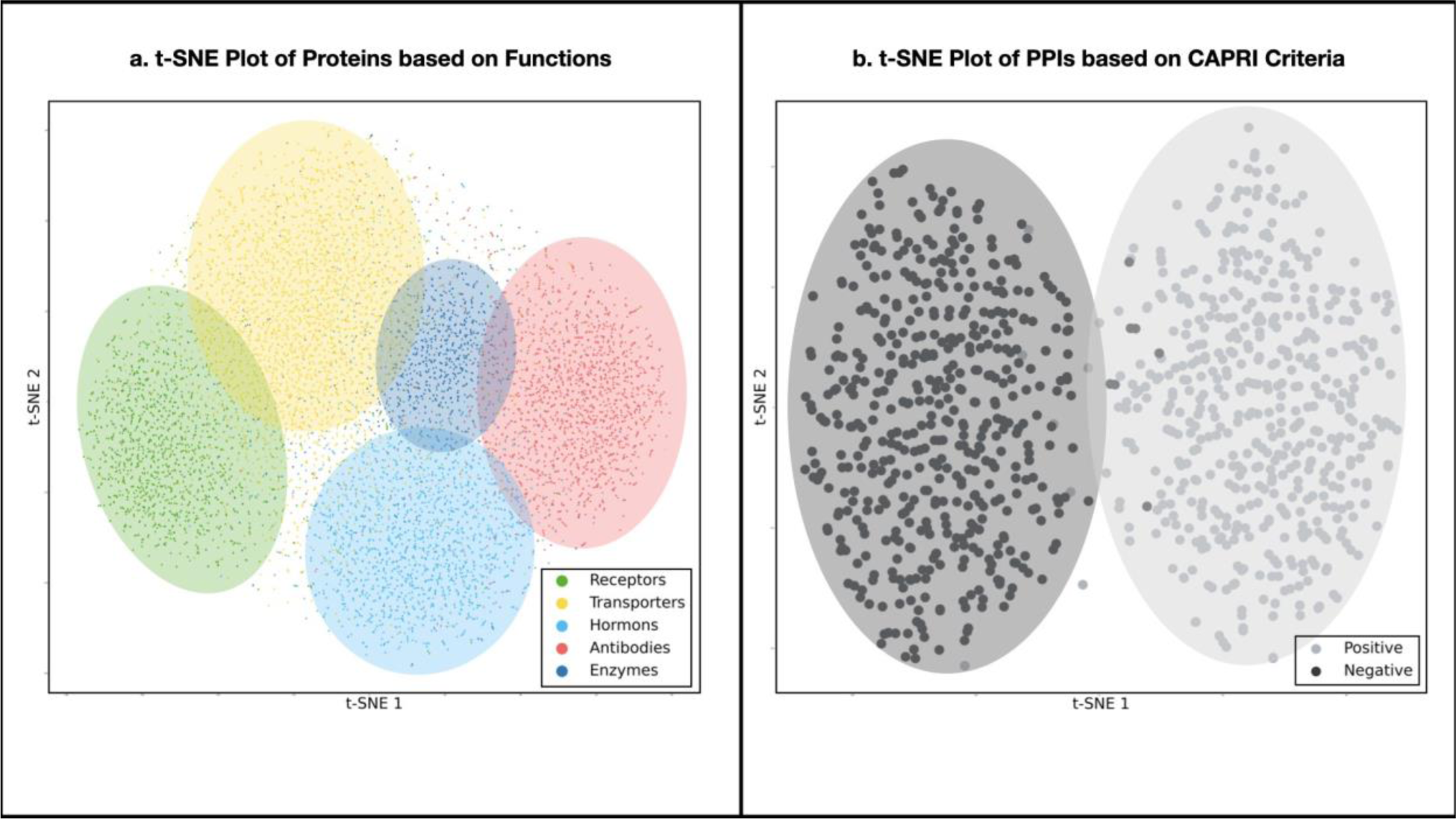
t-SNE graphs of representations learned by ProInterVal. (a) The t-SNE plot represents the clustering of protein functions based on learned representations. Each point corresponds to a protein and is color-coded according to its functional category. (b) The t-SNE plot illustrates the distribution of positive and negative protein-protein interfaces.

In order to conduct a more in-depth analysis of the learned representations of interfaces in our model, a subset of 500 positive and 500 negative interfaces was randomly selected based on CAPRI criteria. The t-SNE graph was then plotted to visualize the distribution and clustering patterns of these interfaces. The results of this analysis are presented in Figure 4.b. As can be seen from the figure, the positive and negative interfaces appear distinctly separated in the embedding space, indicating that our model has effectively learned a robust representation for the validation of protein-protein interactions.

### Ranking of the Positive Interfaces

In order to conduct a comprehensive evaluation of our model, we conducted a ranking analysis to assess its performance in distinguishing a single native interface (positive sample) from a set of approximately 100 incorrect docking models (negative samples). This evaluation involved comparing the performance of our model with GNN-DOVE and DeepRank-GNN, as well as the docking models’ scoring function ZRANK.^35^

We randomly selected 100 positive protein-protein complexes from our DeepInterface test dataset to carry out this analysis. For each of these 100 complexes, we generated 300 decoys using the GRAMM-X protein docking tool.^36^ From the pool of generated decoys, we carefully selected 100 incorrect decoys per complex based on the CAPRI criteria. As a result, we obtain a dataset consisting of 100 incorrect decoys and one native structure per complex.

Next, we applied ProInterVal, ZRANK, GNN-DOVE, and DeepRank-GNN to score and rank these structures. Specifically, we assessed the performance of these scoring tools by examining the position at which the native PDB structure (positive interface) was ranked among the 100 incorrect decoys (negative interface). When we refer to the “top 10%,” it signifies that the near-native interface is positioned within the uppermost 10% of its respective ranked list. Figure 5 indicates the ranking results. Our model achieved a 63% success rate in the top 1% and a 90% success rate in the top 10% and outperformed the others. In practical terms, this indicates that our model placed the native structure among the top 1% for 63 out of the selected 100 complexes and within the top 10% for 90 out of these 100 complexes.

**Figure 5:**
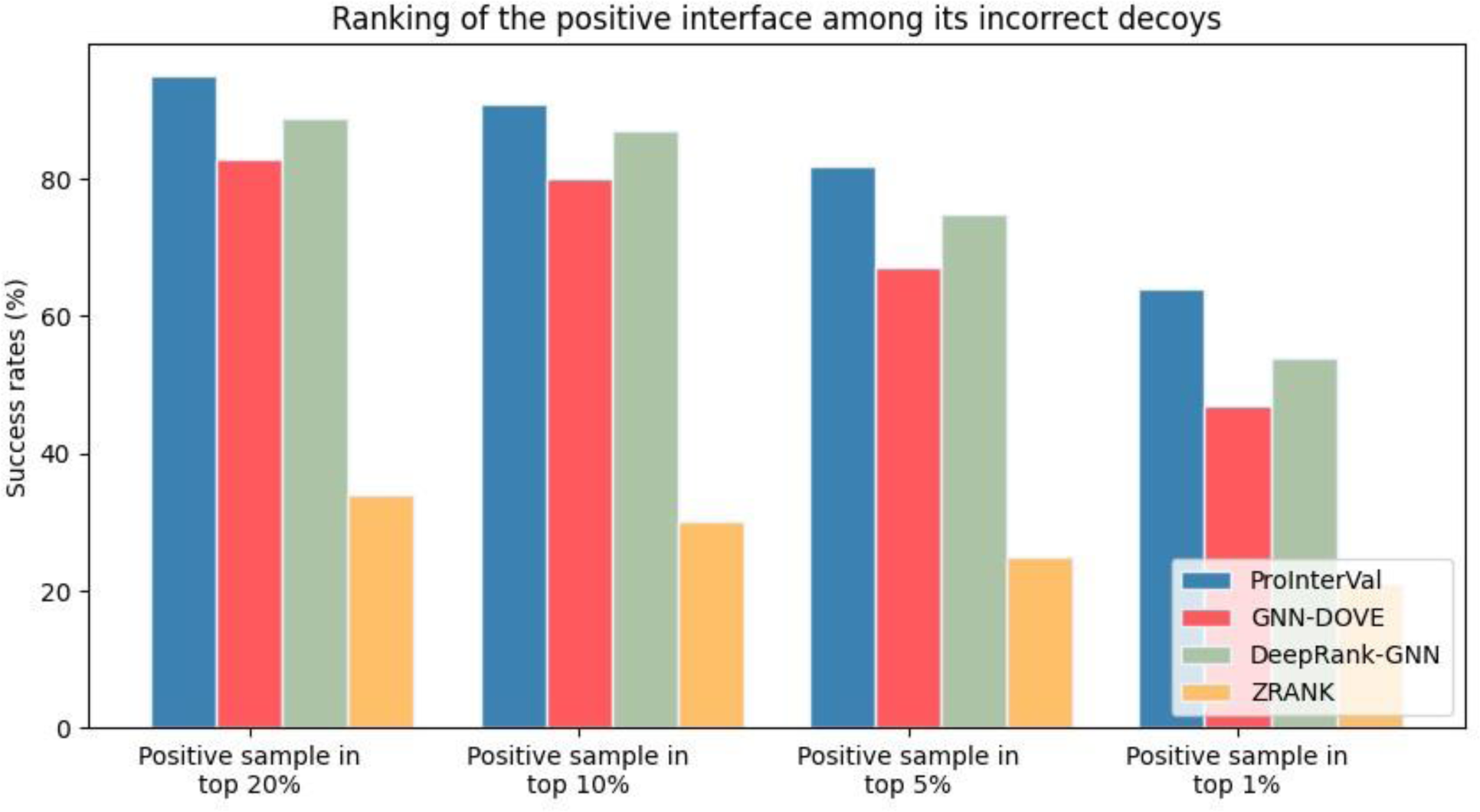
Ranking of the positive interface among the incorrect decoys of the corresponding complex.

The remarkable performance of our model in interface ranking highlights the potential of integrating it into template-based protein-protein interaction (PPI) predictors to improve their overall predictive capabilities.

### Biological vs. Crystal Interfaces Classification

X-ray crystallography has long been the predominant experimental method for determining the 3D structure of proteins, and it remains the most widely used technique as of 2022. While X-ray structures are generally reliable and of high quality, erroneous structures are not uncommon. Common errors include misfitting residues into electron density maps and the presence of synthetic oligomers resulting from crystal packing. The latter can lead to misleading conclusions and impede accurate research findings. Therefore, it is crucial to develop tools to annotate crystallographic dimers as reliable biological or non-biological (crystal) interfaces.

In this section, we assess the performance of ProInterVal in discriminating between biological and crystal interfaces using the DC dataset. Additionally, we compare ProInterVal’s performance with that of DeepRank-GNN.

To train and evaluate ProInterVal, we utilized a dataset comprising 5739 dimers from the MANY dataset. The training set comprised 80% of the dimers, while the remaining 20% constituted the validation set. The network was trained for 50 epochs, and the model with the lowest loss on the validation set was selected as the final model. Subsequently, this model was tested on the DC dataset, which consists of 80 biological interfaces and 81 crystal interfaces with similar interface areas.

ProInterVal achieved approximately 88% accuracy, 88% precision, and 85% F1 score on the test set. The results are shown in the Table 3.

**Table 3:**
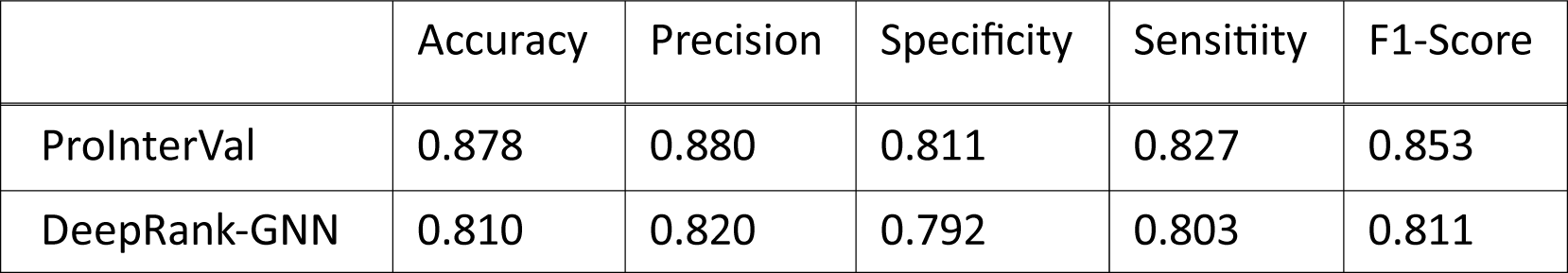
Biological vs. crystal interfaces classification performance evaluation.

|  | Accuracy | Precision | Specificity | Sensitivity | F1-Score |
| --- | --- | --- | --- | --- | --- |
| ProInterVal | 0.878 | 0.880 | 0.811 | 0.827 | 0.853 |
| DeepRank-GNN | 0.810 | 0.820 | 0.792 | 0.803 | 0.811 |

## Conclusion

In this research, we introduced an innovative approach for validating protein-protein interfaces by leveraging learned representations. Our approach utilized a graph-based contrastive autoencoder architecture and a transformer to learn representations of protein-protein interaction interfaces from unlabeled data. This approach allowed us to capture the essential features of the interfaces and encode them into informative representations. The learned representations were then validated using a graph neural network (GNN) model designed for PPI validation.

The results of our experiments on a benchmark dataset demonstrated the effectiveness of our proposed method. We achieved an accuracy of 0.91 for the test set, surpassing the performance of existing GNN-based methods. To our knowledge, GNN-DOVE and DeepRank-GNN are the state-of-the-art GNN-based methods. Therefore, comparing our proposed method to theirs shows the work’s superiority. Our results highlight the potential of learned representations in accurately validating protein-protein interactions.

Furthermore, the learned representations offer versatility in various downstream tasks. For instance, they can be used for the classification of protein complexes as either crystal or biological, helping researchers discern the validity and reliability of crystallographic dimers.

Integrating our model into a template-based PPI predictor holds the potential to enhance the performance of existing prediction methods. By incorporating the learned representations, the template-based predictor can leverage the rich information encoded in the representations, resulting in more accurate and reliable predictions.

As a future work, incorporating sequence information to the structural information can further augment the effectiveness of the representation learning model. By integrating these complementary data sources, the model can capture a more comprehensive view of proteinprotein interactions, considering both the structural characteristics of the interfaces and the sequence patterns involved.

## Data and Software Availability

The information about the data sets and the source code is available on https://github.com/kucosbi/ProInterVal.

## Acknowledgement

All the calculations are performed using the Koc University Advanced Computing Center (KUACC) Facilities. A.G. and O.K. are members of Science Academy, Turkey.

